# Retroactivity induced operating regime transition in a phosphorylation-dephosphorylation cycle

**DOI:** 10.1101/2020.09.30.320606

**Authors:** Akshay Parundekar, Ganesh A Viswanathan

**Affiliations:** Department of Chemical Engineering Indian Institute of Technology Bombay Powai, Mumbai - 400076 India

**Keywords:** Enzymatic cascade, Operating regime, Retroactivity, Rate-balance analysis, Input-output response

## Abstract

Operating regimes characterizing the input-output behaviour of an activated phosphorylation-dephosphorylation reaction cycle (PdPC) such as a single MAPK cascade are dictated by the saturated/unsaturated state of the two underlying enzymatic reactions. Four combinations of the states of these two enzymatic reactions led to identification of distinct operating regimes, *viz*., Hyperbolic (H), Signal transducing (ST), Threshold-hyperbolic (TH) and Ultrasensitive (U). A single PdPC without an explicit feedback have been classically viewed as a module offering signal flow from upstream to downstream, that is, one-way communication. Recently it has been shown that load due to sequestration of the phosphorylated or unphosphorylated form of the substrate by corresponding downstream targets permits retroactive signalling that offers two-way communication. We systematically characterize the operating regimes of a single PdPC subject to retroactivity in either of the substrate forms. We identify five possible regime transitions that could be achieved by increasing the retroactivity strength on either of the two substrate forms. Remarkably, a retroactivity strength of 0.30 in the unphosphorylated form of the substrate is sufficient to induce a transition from ST to H regime indicating that the input-output behaviour of a PdPC is highly sensitive to the presence of a downstream load. Using sensitivity and rate-balance analysis, we show that modulation of the saturation levels of the two enzymatic reactions by increasing retroactivity is the fundamental mechanism governing operating regime transition.

**Highlights:**

- Characterization of operating regimes in the presence of retroactivity.
- Retroactivity can induce a transition between different operating regimes.
- Saturation levels of the two enzymatic reactions govern the regime transition.
- Sensitivity of the protein levels to retroactivity is dictated by saturation levels.

## 1. Introduction

Enzymatic cascades consisting of phosphorylation-dephosphorylation reaction cycles (PdPCs), are crucial, ubiquitously conserved, building-blocks of cellular signalling networks (Widmann et al., 1999; Chang and Karin, 2001). A PdPC employs phosphorylation and dephosphorylation enzymatic reactions, respectively catalysed by kinase and phosphatase, to enable transition of a protein substrate between its two forms, namely inactive and active. PdPCs impart important properties like responsiveness, robustness, specificity, onto a signalling response (Kleiman et al., 2011; Saurin, 2018), weak signal amplification (Blüthgen et al., 2006), signal speed acceleration (Ortega et al., 2002), filter out noise in signal (Thattai and Van Oudenaarden, 2001; Dhananjaneyulu et al., 2012; Baraskar et al., 2013). One such well-known cascade of PdPCs is the Raf/MEK/ERK MAPK cascade, a key signal amplifier and a modulator of prosurvival and pro-apoptotic pathways (Ondrey et al., 1999; Aggarwal et al., 2004; Yang et al., 2016). Aberrant functioning of this cascade has been implicated in many diseases such as cancer (Dhanasekaran and Johnson, 2007; Hanahan and Weinberg, 2011). Detailed understanding of the sustained and transient activation patterns of MAPK cascade can therefore offer useful insights in designing therapeutic strategies for combating certain diseases.

Activation behaviour of a PdPC as a response to a stimulus of certain strength and their dynamic evolution have traditionally been characterized by systematically studying the doseresponse curves permitted by the cascade (Santos et al., 2007; Sepulchre et al., 2012; Frank, 2013). Dose-response curve or the input-output characteristic of the PdPC at steady-state is a map of the abundances of the input kinase and of the active protein (output) of the cascade (Gomez-Uribe et al., 2007). Based on the qualitative nature of the dose-response curve, dictated by the saturated/unsaturated state of the two enzymatic reactions, the activation behaviour of PdPCs have been classified into four distinct operating regimes, *viz*., Hyperbolic (H), Signal transducing (ST), Threshold-hyperbolic (TH), Ultrasensitive (U), each of which display different signal processing capabilities (Goldbeter and Koshland Jr, 1981; Gomez-Uribe et al., 2007). Operating regimes of the MAPK cascades juxtaposed with patient-stratification data have recently been considered in disease prognostics (Fey et al., 2015). Recently, using a hybrid deterministic-stochastic approach for predicting and characterising the input-output behaviour of a single PdP MAPK cycle from ensemble data it was shown that a dose-strength dependent regime transition can occur between H and ST regimes (Parundekar et al., 2019). A quasi-steady-state approximation Michealis-Menten model (Michaelis and Menten, 1913) employed in the hybrid approach (Parundekar et al., 2019) could not explain the observed dose-strength dependent regime transition. A question thus arises as to what could be the mechanism that may govern the observed stimulus-strength dependent operating regime transition under steady-state conditions.

When Raf/MEK/ERK PdPC is modelled without an explicit feedback, signal flow is usually described as a one-way communication, that is, going from upstream to downstream of the cascade (Kholodenko et al., 2010). However, recently, a new type of signalling called retroactive signalling caused by the presence of a downstream load has been considered (Del Vecchio et al., 2008; Ventura et al., 2008, 2010; Catozzi et al., 2016; Grunberg and Del Vecchio, 2020). This phenomenon occurs due to the possibility that the PdPCs are coupled with another downstream cascade/substrate. Either or both forms of the protein involved in a PdPC could be sequestered by another substrate which could be a part of another cascade or simply by a DNA to which one of the forms of the protein is sequestered (Del Vecchio et al., 2008; Ventura et al., 2008, 2009, 2010; Ossareh et al., 2011;). In the case of Raf/MEK/ERK, the sequestration of the phosphorylated ERK could result in a retroactivity in the PdPCs. It has been shown experimentally that retroactivity indeed plays a role in the behaviour of MAPK cascades and other signalling pathways (Ventura et al., 2010; Kim et al., 2010, 2011; Jiang et al., 2011; Jayanthi et al., 2013; Brewster et al., 2014). Presence of retroactivity in a cascade of PdPCs has been suggested to predict a more realistic drug-response curve, that is, an inputoutput behaviour (Ventura et al., 2009).

Inclusion of sequestration effects, which is known to affect the PdPC behaviour (Ventura et al., 2010; Kim et al., 2011) may cause a shift in the operating regimes at deterministic level (Ventura et al., 2009). It is thus likely that incorporating the presence of retroactive signalling might predict the stimulus-strength dependent operating regime transition. In this study, we consider systematically characterising the effect of the presence of substrate or product retroactivity on the operating regimes of MEK/ERK PdPC. Specifically, we show that strength of the retroactive signalling can modulate the nature of the operating regimes and can permit operating regime transitions.

## 2 Mathematical model of a PdPC with retroactive signalling

We consider a single PdPC with retroactive signalling wherein a transition between inactive (*M*) and active (*M_p_*) forms of the protein substrate occurs (Fig. 1). We assume that both forms of the proteins *M* and *M_p_* may be sequestered reversibly, respectively by downstream targets *S_1_* and *S_2_* and thereby incorporate retroactivity in the PdPC (Fig. 1).

**Figure 1:**
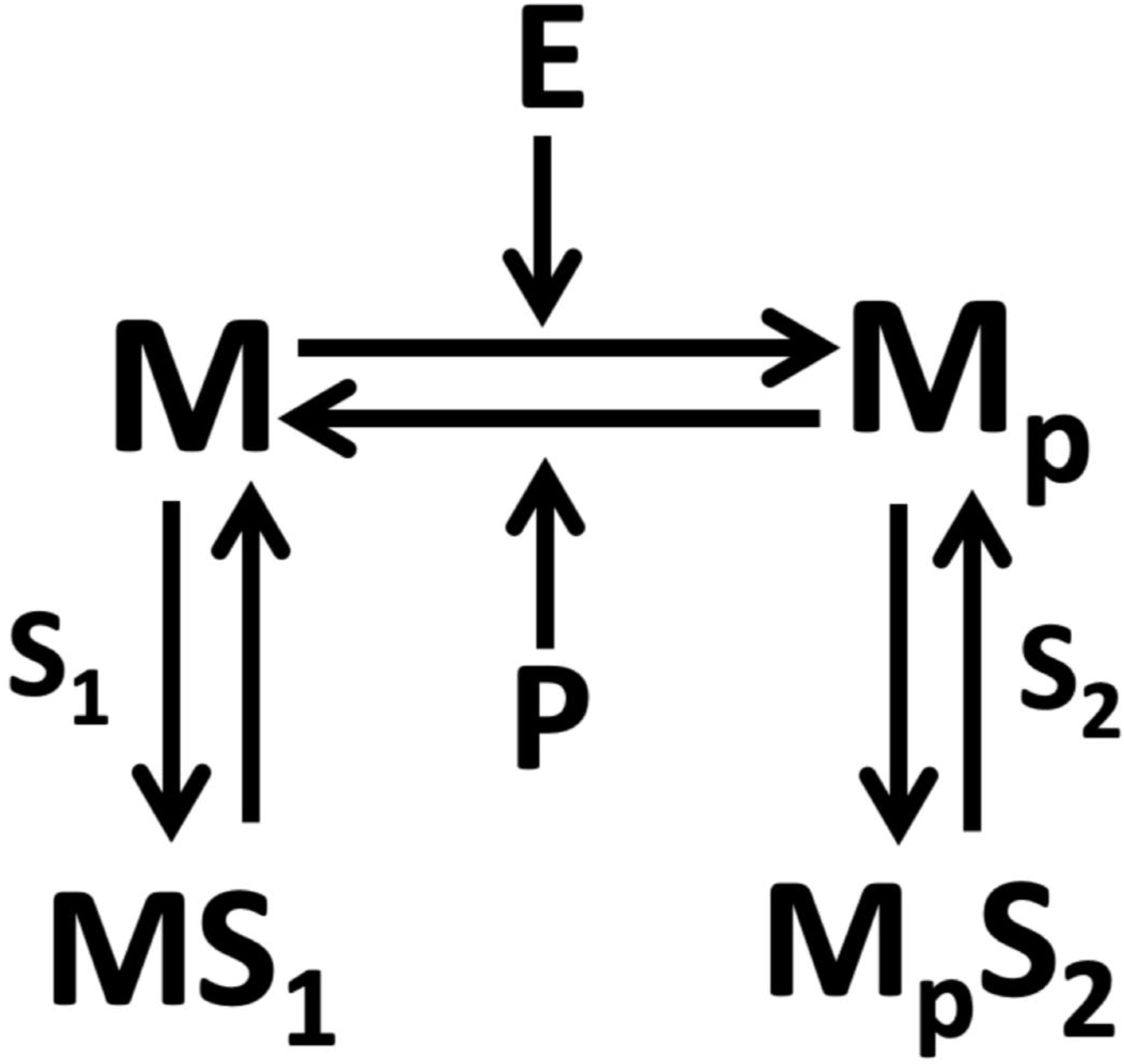
Single PdPC with retroactivity. *M* and *M_p_* are the inactive and active forms of the protein substrate. While *S_1_* and *S_2_* are the downstream targets, respectively of *M* and *M_p_, MS_1_* and *M_p_S_2_* are the corresponding sequestered complexes.

The biochemical reactions corresponding to the PdPC are

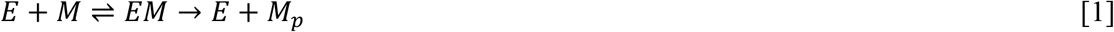

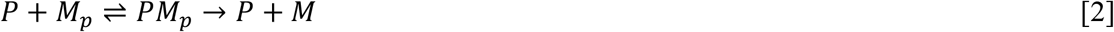

and those capturing the downstream sequestration steps are

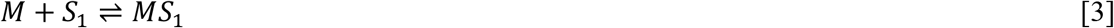

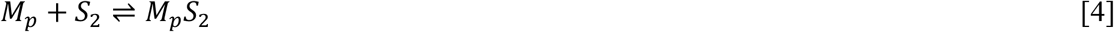

Upon employing pseudo-steady state approximation for the complexes *EM, PM_p_, MS_1_,* and *MpS_2_,* the dynamics of dimensionless concentration of 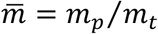, where *m_t_* is the total protein substrate, dictated by the biochemical reactions in Eqs. (1-4) is given by the mathematical kinetic model

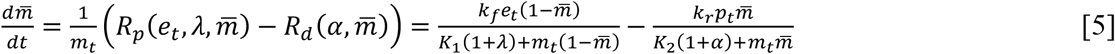

where, *k_f_*, and *k_r_,* respectively are the forward and reverse catalytic rate constants, *e_t_*, and *p_t_*, respectively capture the concentrations of kinase *E,* and phosphatase *P. K*_1_ and *K*_2_ are the Michaelis-Menten (MM) constants for the forward and backward enzymatic reactions, respectively. The retroactivity strengths for sequestration of *M* and *M_p_*, respectively are given by

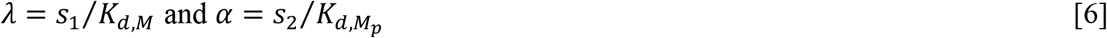

where *s_1_* and *s_2_* are the concentrations of *S_1_* and *S_2_*, respectively. *K_d,M_* and *K_d,M_p__* are the equilibrium constants for binding of *M* and *M_p_* to its respective downstream targets. A detailed derivation of Eq (5) is in Appendix I. Note that the effect of retroactivity of either *M* or *M_p_* or both on the phosphorylation (first term in rhs) and dephosphorylation (second term in rhs) rates in Eq. (5) is quantitatively accounted for by *scaling the MM constants K_1_ and K_2_ with non-zero (positive) values of λ and α, respectively.*

The steady-state solution of Eq. (5) is given by

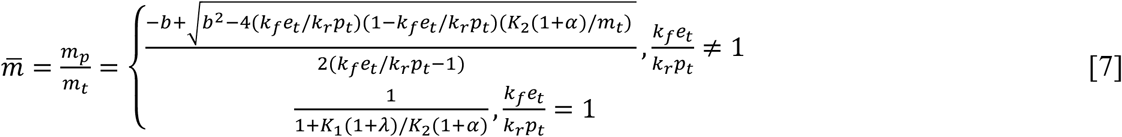

where, *b* = –(*k_f_e_t_*/*k_r_p_t_* – 1) + (*K*_2_(1 + *α*)/*m_t_*)(*k_f_e_t_*/*k_r_p_t_*) + *K*_1_(1 + *λ*)/*m_t_* (Goldbeter and Koshland Jr, 1981; Ventura et al., 2009). Dose-response curve 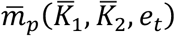 of the PdPC with (or without) retroactivity is essentially the locus of the relationship between 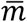 and *e_t_* with all other parameters fixed (Goldbeter and Koshland Jr, 1981; Parundekar et al., 2019). Note that 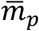 can be drawn using Eq. (7) for (a) without retroactivity by setting *α* = *λ* = 0, (b) with retroactivity only in *M* by setting *α* = 0,*λ* > 0, (c) with retroactivity only in *M_p_* by setting *α* > 0,*λ* = 0, and (d) with retroactivity in both *M* and *M_p_* by setting *α* > 0,*λ* > 0 (Ventura et al., 2009). Since introduction of retroactivity tantamount to scaling of the MM constants (Eq. 5), for the sake of brevity, we define effective MM constants 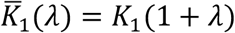 and 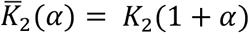, which when *λ* or *α* set to zero will correspond to the case absence of retroactivity in *M* or *M_p_*, respectively. Dose-response curve 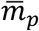 can be classified into four distinct operating regimes by contrasting it with 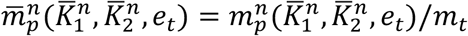, where superscript *n*=H, ST, TH, U indicates regime-specific nominal profile specified by 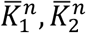 in Table 1 (Methods). As a reference, we employ the nominal profiles corresponding to the case wherein retroactivity is absent. This classification has been suggested by Gomez-Uribe (Gomez-Uribe et al., 2007) and adopted in several recent studies (Straube, 2017; Parundekar et al., 2019).

**Table 1:** Michealis-Menten constants 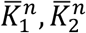 used to arrive at the nominal profiles of the four regimes (Gomez-Uribe et al., 2007; Parundekar et al., 2019).

| Regime | ST | H | TH | U |
| --- | --- | --- | --- | --- |
| $\bar{K}_1^n, \bar{K}_2^n$ | 10,10000 | 10000,10000 | 10000,10 | 10,10 |

## 3. Results

### 3.1 Retroactivity impacts operating regimes

In order to study the effect of retroactivity on the dose-response curve, we adopt the same strategy prescribed by Gomez-Uribe (Gomez-Uribe et al., 2007) to characterize the operating regimes in the presence of a downstream load on *M* or *M*_p_. We limit the scope of this study to the presence of retroactivity in either *M* or *M_p_*. (Systematic characterization reported here, without loss of generalization, can be used for the case where retroactivity may be present in both *M* and *M_p_*, simultaneously.) Unless otherwise explicitly stated, for the rest of study, parameters assumed are *k_r_* = *k_f_* = 0.01s^-1^, *p_t_* = 200*nM* and *m_t_* = 1000*nM* (Gomez-Uribe et al., 2007; Parundekar et al., 2019).

In order to assess if retroactivity impacts the nature of the operating regime for a certain set of parameters, we consider a dose-response curve in the ultrasensitive (U) regime in the absence of retroactivity (*α* = *λ* = 0), when 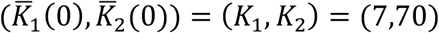. Figure 2A shows this dose-response curve (solid yellow) contrasted against the nominal profile for U regime (dashed blue) used for identifying the regime to which it belongs to. Introduction of retroactivity in *M_p_* with a strength of *α* = 27 (and *λ* = 0) resulting in 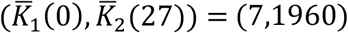 causes shifting of the dose-response curve (solid purple curve in Fig. 2A) to the left. 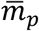 for case of *α* = 27 belongs to the signal transducing (ST) regime indicating the possibility of retroactivity induced transition of operating regimes. (For the sake of comparison, we present 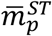 (dashed red curve) in Fig. 2A.). We further show that introduction of (a) retroactivity in *M_p_* can induce regime transition from hyperbolic (H) at 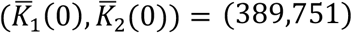 to ST regime at 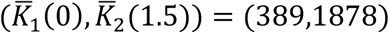 (Fig. 2B) and (b) retroactivity in *M* can induce operating regime transition from ST to H (Suppl Mat., Text S1). Given that the presence of a downstream load can cause a regime shift, we ask a question as to what are the other possible transitions in the presence of retroactivity. The primary goal of this study is to systematically understand the effect of retroactivity in *M* or *M_p_* on the operating regimes.

**Figure 2:**
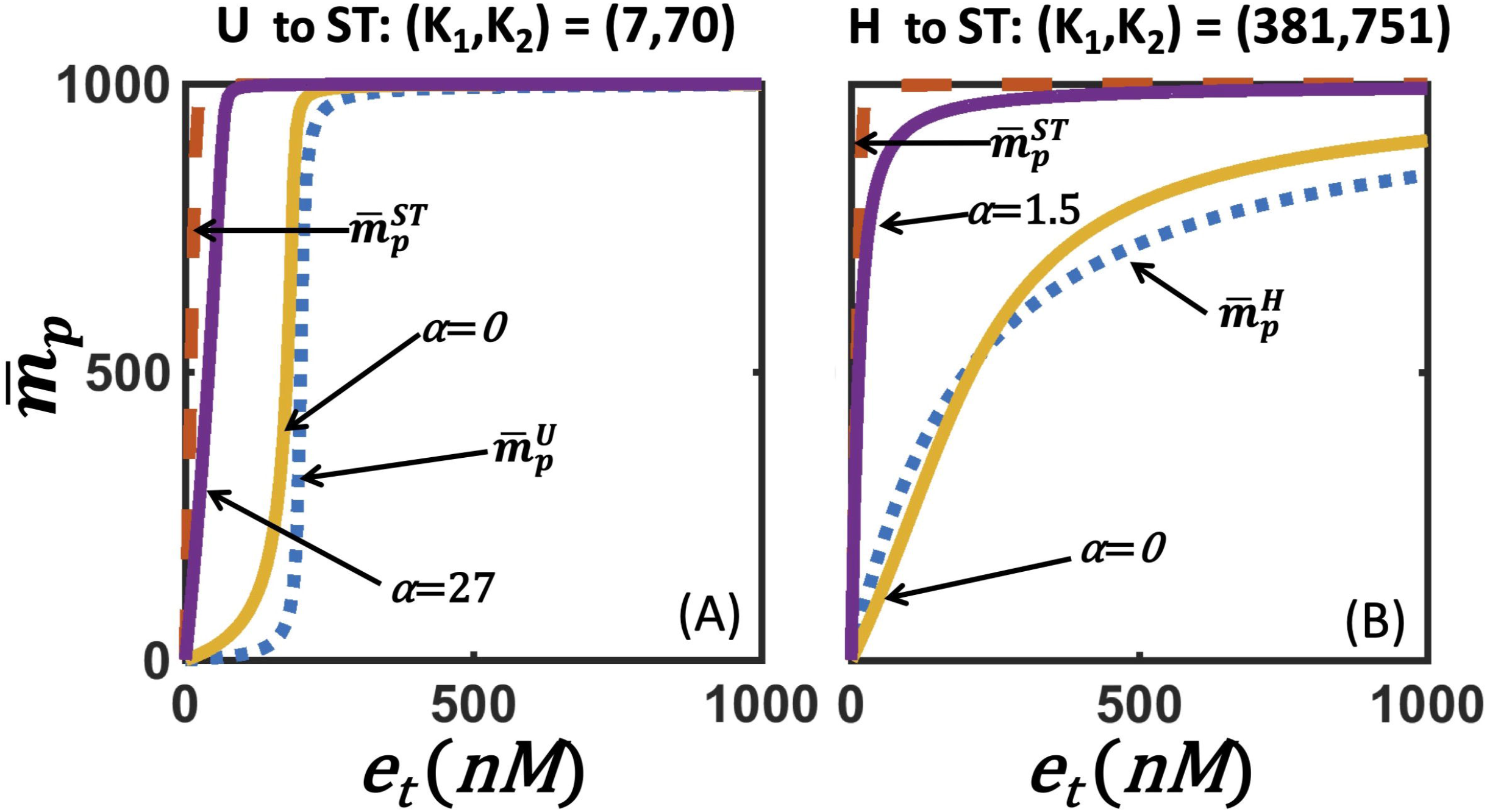
Retroactivity in *M_p_* inducing operating regime transition from (a) U at 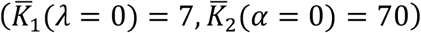 to ST at 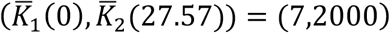 and (b) H at 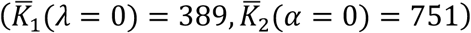 to ST at 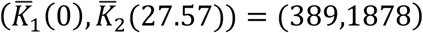.

### 3.2 Retroactivity strength dictates nature of regime transition

Retroactivity introduces a scaling for the Michaelis-Menten constants (Eq. 5) and thereby affects the steady-state behaviour (Eq. 7). As a result, in order to study the effect of retroactivity strength on the operating regimes, it is sufficient to understand how the parameter space of effective Michaelis-Menten constants 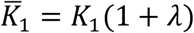 and 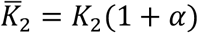 is partitioned into different input-output behaviours. Note that replacing *K*_1_ (1 + *λ*) and *K*_2_(1 + *α*) in Eq. (7) respectively with 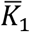 and 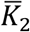 makes the steady-state solution form for the case when retroactivity is present is similar to that of an isolated PdPC. Thus, knowledge of the boundaries of the different operating regimes in the planes of 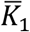 and 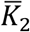 could be directly used to decipher the effect of retroactivity on the dose-response curves exhibiting a certain input-output characteristic by varying *λ* or *α*.

Next, we implemented an optimization problem to delineate the parameter space 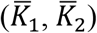 corresponding to the four distinct operating regimes. For the ease of constructing the map, assuming *α* = *λ* = 0, for an operating regime, say H, after specifying a 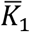, we identified 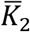 by increasing retroactivity strength *α* such that the candidate doseresponse curve 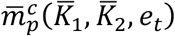 satisfied the relative distance criterion

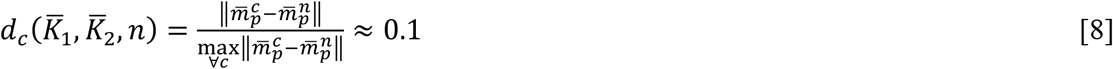

for all *n*=H, ST, TH, and U. This criterion is based on the metric suggested by Gomez-Uribe et al. (Gomez-Uribe et al., 2007) and recently used in Parundekar et al. (Parundekar et al., 2019). Note that for every candidate dose-response curve there will be four distances, each corresponding to a comparison with four regime-specific nominal profiles. Achieving this objectively requires estimation of 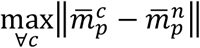 in Eq. (8) *a priori.* However, the information about regime to which 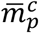 belongs to is unavailable. In order to address this, we first created a randomly chosen parameter-profile database containing 140000 sets of 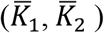 chosen using stratified random sampling (Methods) across five orders of magnitude range each tagged to its dose-response curve 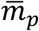. (Note that the maximum possible value that an element in 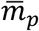 can take is 1 (Parundekar et al., 2019).) Next, we performed an optimization (Methods) for finding 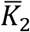 that satisfies Eq. (8) and its corresponding 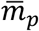. In the five-orders of magnitude range considered, finding 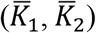 whose corresponding 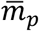 satisfied Eq. 8 upon choosing *n*=H enabled identifying the boundary for the H regime in the planes of effective Michaelis-Menten constants (orange lines in Fig. 3). Note that the dashed lines correspond to those 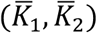 on the boundary sourced directly from the database. We repeated the entire procedure to find the boundaries corresponding to U (blue), ST (yellow), and TH (red) regimes (Fig. 3). We note that upper boundary of the ST regime is an exception. While constructing the upper boundary for ST regime, we observed that the dose-response curve is insensitive to 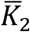 beyond a certain limit after which 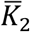 has no effect on 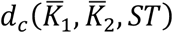. Therefore, for representation purposes, we fixed the upper boundary for ST (yellow) at 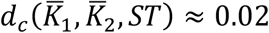 by accordingly modifying Eq. 8. Note that as a direct consequence the dose-response curves well beyond the upper boundary of ST will belong to the signal-transducing regime. Metric adopted in Eq. 8 by and large separates the regions where these four regimes exist, as has also been reported in Gomez-Uribe et al. (Gomez-Uribe et al., 2007).

**Figure 3:**
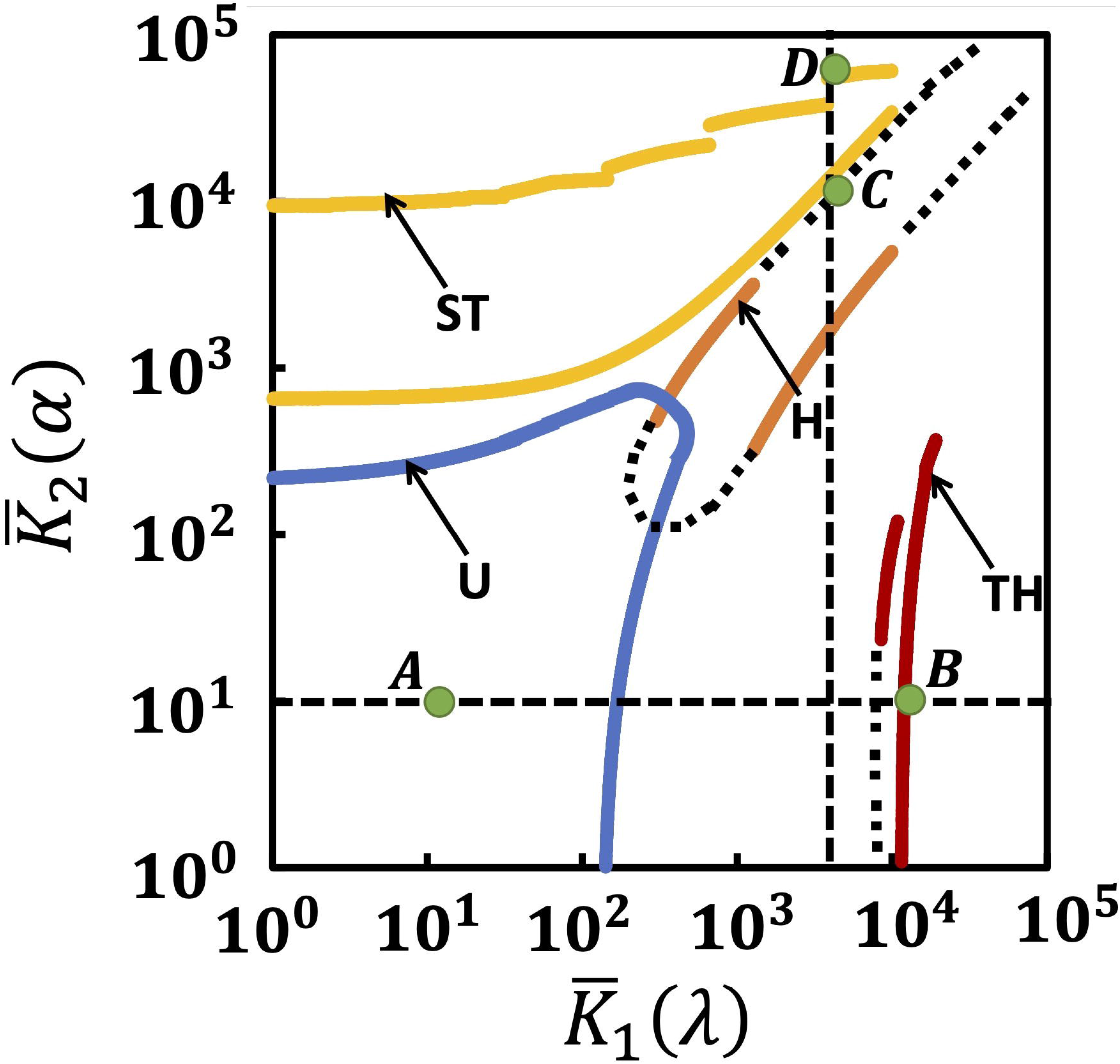
Boundaries of the four operating regimes (a) hyperbolic (H, blue), (b) signal transducing (ST, green), (c) threshold-hyperbolic (TH, red), and (d) ultrasensitive (U, yellow) in the planes of 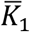 and 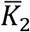 for *λ* = *α* = 0. Dashed lines correspond to those 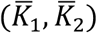 sourced directly from the database.

Since changing retroactivity strength can independently modulate the Michaelis-Menten constants, manipulating 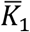, or 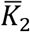 or both could cause a shift in the characteristic input-output behavior. Specifically, by increasing the strength of the load in *M* causing change in 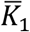, while keeping 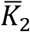 constant, a dose-response curve in U or ST, respectively can shift to TH or H. For example, 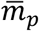(10,10) in U regime (point A in Fig. 3) would shift to TH (point B in Fig. 3) upon increasing 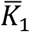, to 10000. Similarly, while maintaining 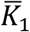 constant, an increase in the retroactivity strength in *M_p_* leading to proportional change in 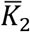 could lead to four possible regime transitions, *viz*., U to ST, TH to H or ST, and H to ST. For example, 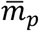 (6000,6000) in H regime (point C in Fig. 3) transitions into ST regime (point D in Fig. 3) when 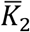 is scaled to 60000. For a given source profile specified by a certain 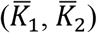 with no retroactivity either in *M* or *M_p_*, while maintaining 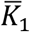, or 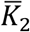 constant, the minimum load *α_min_* or *λ_min_*, respectively required for inducing a regime transition is sensitive to the chosen 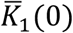 or 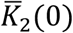 (Suppl Mat, Text S2). The sensitivity analysis showed that *λ_min_* = 0.3 is sufficient to induce ST to H transition due to retroactive signalling.

### 3.3 Saturation level of the two enzymatic reactions governs the retroactivity induced regime transition

Since the dose-response curve 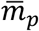 explicitly depends on 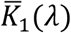 and 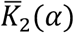 (Eq. 7), understanding how load strength *λ* or *α* influences the input-output behaviour may offer useful insights into what causes retroactivity driven operating regime transition. In order to assess the regime-specific impact of retroactivity on the input-output behaviour, we systematically analyse the sensitivity of the dose-response curves with respect to retroactivity strengths *λ* and *α*.

The sensitivity of 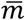 with respect to retroactivity strength *λ* and *α*, respectively are quantitatively captured by

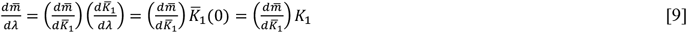

and

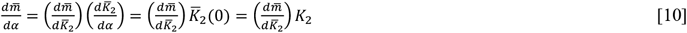

for a finite downstream load. Detailed expressions of these are in Appendix II. Eqs 9 and 10 show that the presence of retroactivity in *M* or *M_p_* introduces a constant scaling of 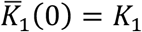 or 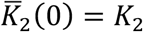, respectively to the sensitivity with respect to the corresponding Michealis-Menten constant. We present the sensitivity effects for two possible transitions by first fixing 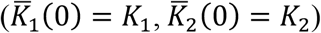 in U-regime with no retroactivity and then increasing the retroactivity strength.

#### 3.3.1 U to TH transition due to retroactivity in M

In Fig. 4A, for 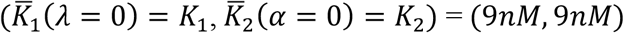 with *d_t_*(9*nM*, 9*nM, U*) = 0.013, maintaining *α* = 0, we show the modulation of sensitivity by *λ* causing proportional scaling of 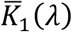 and by total upstream enzyme concentration *e_t_*. For a given *λ*, a peak in the sensitivity occurs. An increase in *λ* in 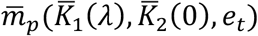 leads to a decreased negative sensitivity for all values of *e_t_* and also causes a shift in the peak location away from the ordinate. This shift is correlated to the corresponding increase in the EC50, that is, the enzyme concentration at which 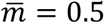, as has been reported in Ventura et al. (Ventura et al., 2009). This behaviour is strongly dictated by the steady-state levels of *M_p_* that the solution (Eq. 7) permits with increasing *λ*. Since the steady-state level is a balance between the two terms in rhs in Eq. 6, insights into the effect of *λ* on 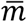 can be deciphered from the phosphorylation and dephosphorylation rate curves. In Fig. 4B, we present the rate-balance plot consisting of the locus of 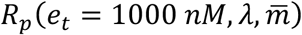 and 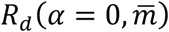, intersection of which for a specific *λ* gives the corresponding steady-state level. Note that in the U regime, 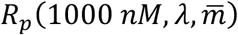 (Fig. 4B, orange) and 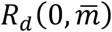, (Fig. 4B, dashed) curves, respectively correspond to the case where both enzymatic reactions are close to saturated, that is, all of the enzymes are bound to its substrate. For the chosen *e_t_*, at *λ* = 0, the intersection occurs in the region where *R_p_* fall rapidly and *R_d_* levels-off, leading to 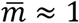. Increase in *λ* leading to a proportional increase in 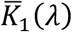 forces the *R_p_* curve to move from saturated to unsaturated behavior (Fig. 4B). Depending on the extent of unsaturation introduced in the phosphorylation reaction, shift in the intersection point of the rate curves occurs either in the region where *R_d_* curve levels-off to maximal rate (Fig. 4B, *λ* = 0 to 400), in the intermediate 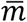 range, or in the range where the *R_d_* curve falls rapidly. This suggests that the effect of *λ* on the sensitivity is pronounced when the intersection of the two rate curves occurs in the region where the *R_d_* curve levels-off to a maximum rate. The effect continues to reduce drastically with further increase in *λ* wherein the intersection falls in the region where *R_d_* falls rapidly (Fig. 4B, dashed line and Fig. 4A) and thereby pushing the steady-state levels lower. For the set of parameters considered, a minimum *λ* of 960 is needed to make the phosphorylation reaction sufficiently unsaturated that the dose-response curves transition into the TH regime (dashed line in Fig. 4A and 4C). The dose-response curve at the transition is at a relative distance of 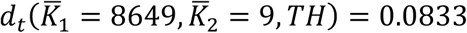 from the TH nominal profile 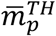

**Figure 4:**
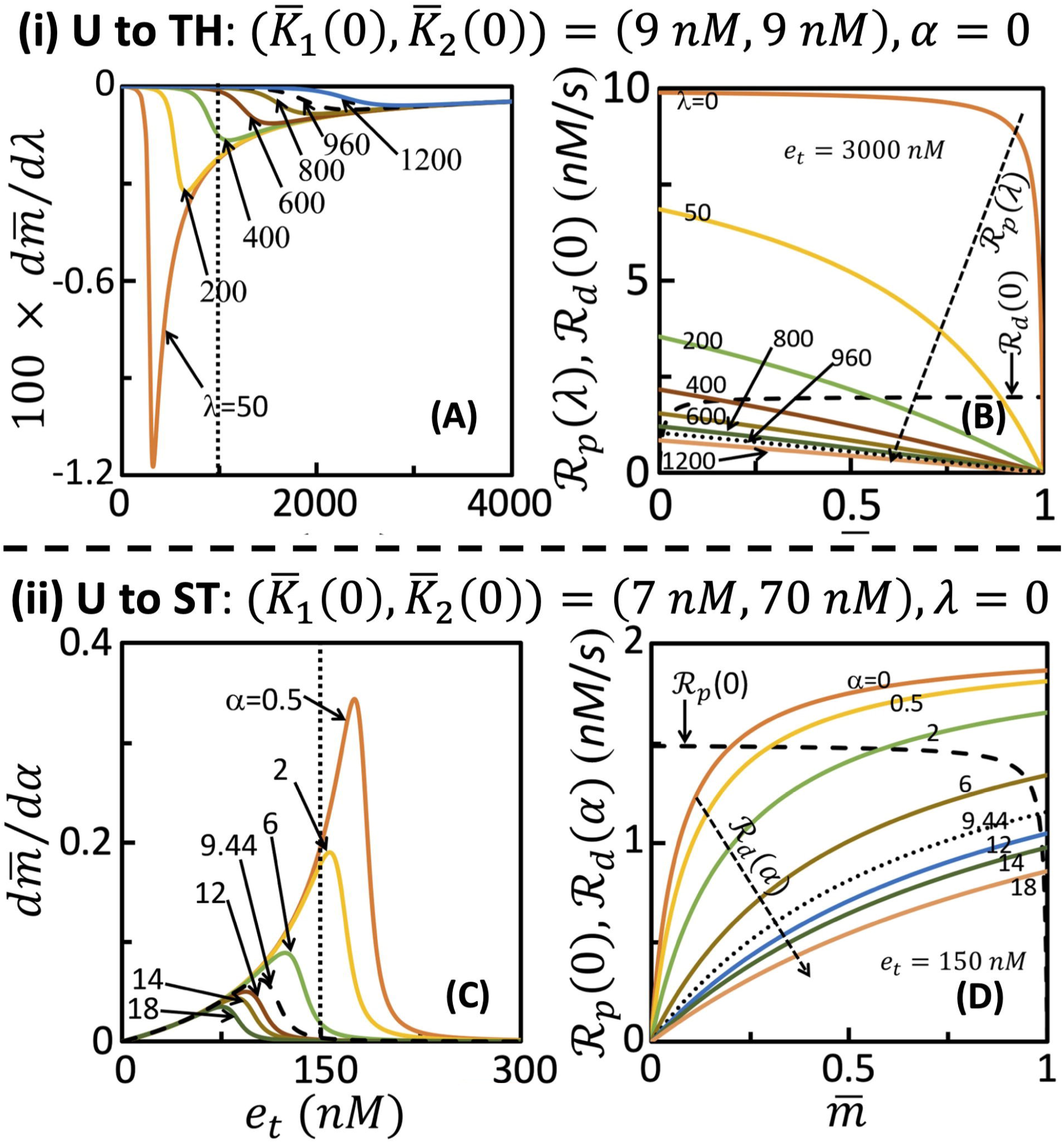
Effect of retroactivity strength on sensitivity of steady-state level with respect to (a) load on *M* when a = 0 and (c) load on *M_p_* when *λ* = 0. Rate-balance plots showing retroactivity strength *λ* and *α*, respectively introducing unsaturation in (b) phosphorylation reaction for *e_t_* = 1000*nM* and (d) dephosphorylation reaction for *e_t_* = 150*nM*.

#### 3.3.2 U to ST transition due to retroactivity in M_p_

For 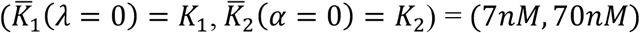 with *d_t_*(7 *nM*, 70 *nM, U*) = 0.049, Fig. (4C) shows that an increase in the retroactivity strength *α* in 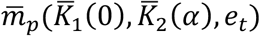 while maintaining *λ* = 0 causes (a) reduction in the (positive) sensitivity at all enzyme concentrations *e_t_* and (b) increase in ‘*e_t_*’ at which the peak occurs. A rate-balance analysis at *e_t_* = 1000*nM* shows that when *α* = 0, the intersection of the two rate-curves occurs in the region where *R_p_* levels-off at a certain maximum rate (Fig. 4D). An increase in *α*, that is, 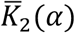 leads to a shift of the *R_d_* curve from saturation to unsaturation. The extent of unsaturation introduced is modulated by increasing a and thereby causing a shift in the intersection of the two curves away from the ordinate leading to a higher steady-state level (Fig. 4D). As a result, similar to the case considered earlier (in section 3.3.1), the sensitivity is more pronounced in the a range where the steady-state concentration achieved is due to intersection of the rate-curves in the region where *R_p_* levels-off at a maximal rate (Fig. 4D, dashed and Fig. 4C). Minimum *α* = 9.44 is necessary for the dose-response curves thus obtained to transition into ST regime having a 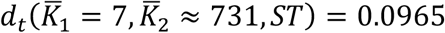.

The effect of retroactivity strength on the sensitivity for other three regimes are in Suppl Mat., Text S3. While the mechanism governing ST to H transition (Suppl Mat., Fig. S3) due to load in *M* mimics that for U to TH transition (section 3.3.1), TH to H transition (Suppl Mat., Fig S4) due to load in *M_p_* is dictated by the underlying principle observed for U to ST transition (section 3.3.2). These observations suggest that the four transitions between different operating regimes can be facilitated by varying the strength of retroactivity that may be present in *M* or *M_p_*. However, for the case of H to ST transition, the two enzymatic reactions continue to remain unsaturated before and after the regime transition (Suppl Mat., Fig S5).

## 4. Discussion and Conclusion

Input-output behaviour of an activated phosphorylation-dephosphorylation cycle (PdPC) has been studied extensively due to its ability to orchestrate cell fate in direct and indirect contextdependent manner (Chang and Karin, 2001; Dhanasekaran and Johnson, 2007; Parundekar et al., 2019). Michealis-Menten constants (MM) dictated saturated/unsaturated state of the two enzymatic reactions facilitates placing steady-state dose-response curves of a PdPC into Signal transducing (ST), Hyperbolic (H), Threshold hyperbolic (TH) and Ultrasensitive (U) operating regimes (Gomez-Uribe et al., 2007). The unphosphorylated (*M*) and phosphorylated (*M_p_*) forms of the protein substrate involved in the PdPC can be sequestrated by respective downstream targets. The sequestration dictated load or retroactivity on the upstream protein levels introduces a two-way signal flow permitting modulation of the steady-state behaviour (Del Vecchio et al., 2008; Ventura et al., 2010). In this study, we systematically show that the presence of retroactivity in *M* or *M_p_* can shift the input-output behaviour from one operating regime to another by modulating the level of saturation of the enzymatic reactions. In particular, we demonstrate five possible transitions: (a) U to TH and ST to H caused by retroactivity in *M* and (b) U to ST, TH to H, and H to ST by that in *M_p_*. Our study corroborates the recent experimental observations that a stimulus-strength dependent shift in the operating regimes is possible in a single MAPK cascade (Ventura et al., 2010, Parundekar et al. 2019).

Using a pseudo-steady state approximated model of the PdPC, we systematically identified the MM constants range that permit four distinct operating regimes in the presence of retroactive signalling (Fig. 3). Interestingly, simulations revealed that presence of a very small load can cause a shift in the operating regime. For instance, a retroactivity strength of 0.3 can cause the dose-response curve at 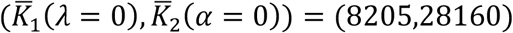 belong to ST regime to transition into H regime. Several downstream targets that could sequester proteins in MAPK cascades have been reported (Kim et al., 2011). Thus, while analysing such a behaviour in PdPC using experimental data, ignoring the hidden retroactive signalling effect, however small, can lead to incorrect prediction of the underlying operating regime.

While in this study, we only considered increasing the retroactivity strength to trigger a regime transition, in principle, if downstream sequestration were already present, its strength can be decreased too. Decreasing the retroactivity strength can predict five transitions that are essentially the reverse of those reported in this study. Further, simultaneous increase (decrease) of the retroactivity strengths in *M* and *M_p_* can lead to a transition from U to H (H to U). Thus, the operating regime boundaries reported in Fig. 3 permits prediction of all 12 possible regime transitions. We further note that the U and H regimes have a slight overlap in the 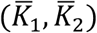 space, as has been reported by Gomez et al. (Gomez-Uribe et al., 2007).

Using sensitivity and rate-balance analysis, we demonstrated that modulation of the saturation levels of the two enzymatic reactions by increasing the retroactivity strength is the fundamental reason for the operating regime transition observed in all cases except H to ST transition. In particular, we show that increase in retroactivity (a) in *M* leads to increasing unsaturation in the phosphorylation reaction and (b) in *M_p_* makes dephosphoryation reaction more unsaturated (Fig. 4). This is due to the fact that the steady-state level of *M_p_*, the active form, is sensitive to changes in the retroactivity strength. While increasing the strength of retroactivity in *M* causes a decrease in the (negative) sensitivity of the steady-state level, that in *M_p_* leads to marked reduction in the (positive) sensitivity. This sensitivity to retroactive signalling can be capitalized upon to modulate the nature of response of PdPC. Synthetic biology tools are becoming available for tweaking the binding sites of targets to which the protein substrate, active/inactive forms may bind and thereby enabling control of the extent of sequestration (Morsut et al., 2016). The nature of sensitivity effect that retroactive signalling bestows on the steady-state levels demonstrated in this study can be of immense value for precise engineering of a cell to control and modulate the input-output behaviour.

## 5. Methods

### 5.1 Regime identification

The regime that a dose-response curve 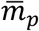 belongs to is identified by contrasting it with the four nominal profiles 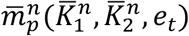 where superscript *n*=H, ST, TH, U. The dose-response curve 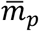 is tagged to below to a certain regime H, ST, TH, or U if the relative distance between 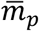 and the corresponding nominal profile is within 10%.

### 5.2 Stratified random sampling

The two stratification cut-off points were chosen is such a way that (a) 60000 samples were chosen in the (*K*_1_, *K*_2_) range of [0-1600, 0-1600], and (b) 10000 samples each were chosen in the range [0-50, 0-10000] and [0-10000, 0-50]. In both these cases uniform distribution was used for sampling. Samples were chosen in such a way that those representing phosphorylation or dephosphorylation or both reaction(s) being saturated was at least 10% greater than those for unsaturated in both the enzymatic reactions of the PdPC.

### 5.3 Optimization for finding operating regime boundaries

The boundary for a specific regime was obtained by seeking 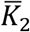 that satisfied the objective function in Eq. 8 which was solved using nonlinear optimization function *“fmincon”* implemented in Matlab® (Mathworks, 2017). A tolerance of 1e^-6^ was set as convergence criteria to the optimization problem. Optimizer convergence was sluggish in the presence of steep gradients and in these cases, the 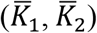 samples from the database that satisfied Eq. 8 was used.

## Acknowledgements

We thank Department of Science and Technology, Government of India for funding this study.

## Appendix I: Derivation of the mathematical kinetic model

The dynamics of the sequestration reactions in Eqs (3-4) are given by

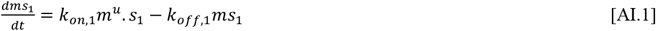

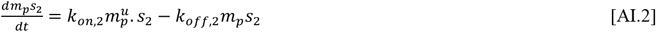

where, *m^u^* and 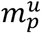, respectively are concentrations of *M*and *M_p_* not bound to their downstream targets. We assume same binding constant values for the two sequestration events, that is, *k*_*on*,1_ = *k*_*on*,2_ and *k*_*off*,1_ = *k*_*off*,2_ (Ventura et al., 2009). Using the quasi-steady state approximation (QSSA) for Eqs (AI.1 and AI.2), the retroactivity strengths on *M* and *M_p_* are *λ* = *ms*_1_/*m^u^* = *s*_1_/*K_d,m_* and 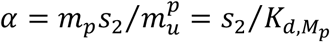, where *K_d,M_* = *K_d,M_p__* = *k*_*off*,1_/*k*_on,1_ The conservation relations for total concentrations of *M*, *M_p_*, substrate, *E* and *P*, respectively are *m* = *m^u^* + *ms*_1_ = *m^u^*(1 + *λ*), 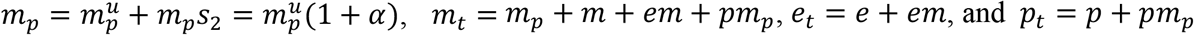.

We next assume QSSA for the dynamic model of biochemical reactions Eqns (1, 2)

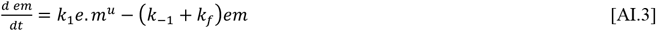

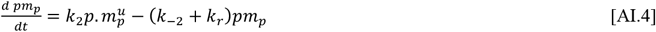

where *k_1_* and *k_−1_* are the forward and backward rate constants for the biochemical reaction in Eq. (1) and *k_−1_* and *k_−2_* are those for the reaction in Eq. (2). Incorporating the conservation relations, the concentration of the intermediate complexes *EM* and *PM_p_*, respectively are *em* = *e_t_.m*/(*K*_1_ + *λ*) + *m*) and *pm_p_* = *p_t_.m_p_*/*K*_2_(1 + *α*) + *m_p_*, where, the Michaelis-Menten constants *K*_1_ = (*k*_-1_ + *k_f_*)/*k*_1_ and *K*_2_ = (*k*_-2_ + *k_r_*)/*k*_2_. Incorporating expressions for *em* and *pm_p_* into dynamics of total phosphorylated species *m_p_*

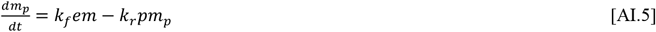

results in Eq. (5).

## Appendix II: Sensitivity of steady-state level to retroactivity strength

Assuming 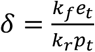 the expressions for sensitivity of the steady-state level to retroactivity strength due to sequestration of *M* by a downstream target (Eq. 9) is

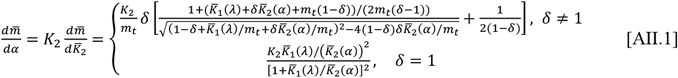

and due to sequestration of *M_p_* by its downstream target (Eq. 10) is given by

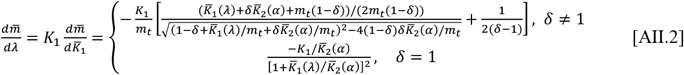

where, 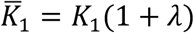 and 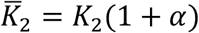. Definition of all other quantities are as that in Eq. 5

## Supplementary Material

**Text S1:** Dose-response transition from Signal-Transducing to Hyperbolic regime induced by substrate retroactivity.

**Text S2:** Minimum retroactivity strength required to transition from one regime to another.

## Footnotes

1 Johnson, K.A., Goody, R.S., 2011. The original Michaelis constant: translation of the 1913 Michaelis–Menten paper. Biochemistry 50, 8264–8269. https://doi.org/10.1021/bi201284u

## Notes

### Competing Interest Statement

The authors have declared no competing interest.

