## Supplementary text for "Retroactivity induced operating regime transition in a phosphorylation-dephosphorylation cycle"

**S1. Dose-response transition from Signal-Transducing to Hyperbolic regime induced by substrate retroactivity.**

We illustrate here the impact of substrate retroactivity in inducing transition in the nature of operating regime from Signal-transducing (ST) to Hyperbolic (H). In Fig S1, we show that $\bar{m}_p(9{,}2000)$ in ST regime transitions into H regime when $\lambda = 446$ leading to $\bar{K}_1 = 4023$ while maintaining $\alpha = 0$.

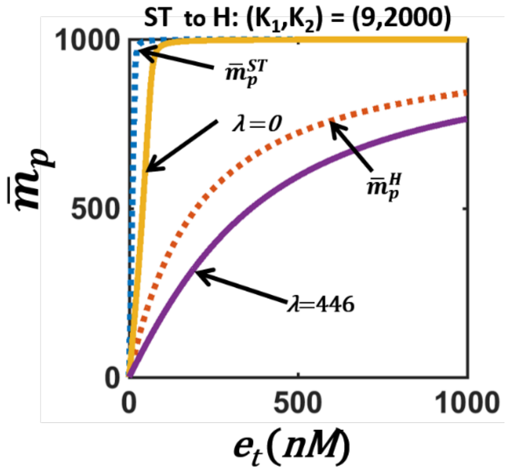


**Figure S1**: Transition of dose-response curve from ST to H

**S2. Minimum retroactivity strength required to transition from one regime to another.**

In order to find the minimum retroactivity strength needed to permit a regime transition, we consider each of the five possible transitions achieved by increasing the load $\lambda$ or $\alpha$. We demonstrate by considering the case of ST to H transition. For every $(\bar{K}_1(\lambda = 0), \bar{K}_2(\alpha = 0))$ on the ST regime boundary, we estimate $\lambda_{min}$, the minimum load on $M$ needed to proportionally scale $\bar{K}_1(\lambda_{min})$, while keeping $\bar{K}_2(\alpha = 0)$ constant, to cause the dose-response curve (solution of Eq. 5 (main text)) to belong to the H regime as per the condition in Eq. 8 (main text). Since this minimum load is sensitive to the chosen $\bar{K}_2(\alpha = 0)$, we find $\lambda_{min}$ for all $\bar{K}_2(\alpha = 0)$ considered along the ST regime boundary. This procedure is repeated the remaining four regime transitions. The sensitivity to the minimum load $\lambda_{min}$ or $\alpha_{min}$ due to $\bar{K}_2(\alpha = 0)$ or $\bar{K}_1(\lambda = 0)$ for the considered five transitions are presented in Fig S2.

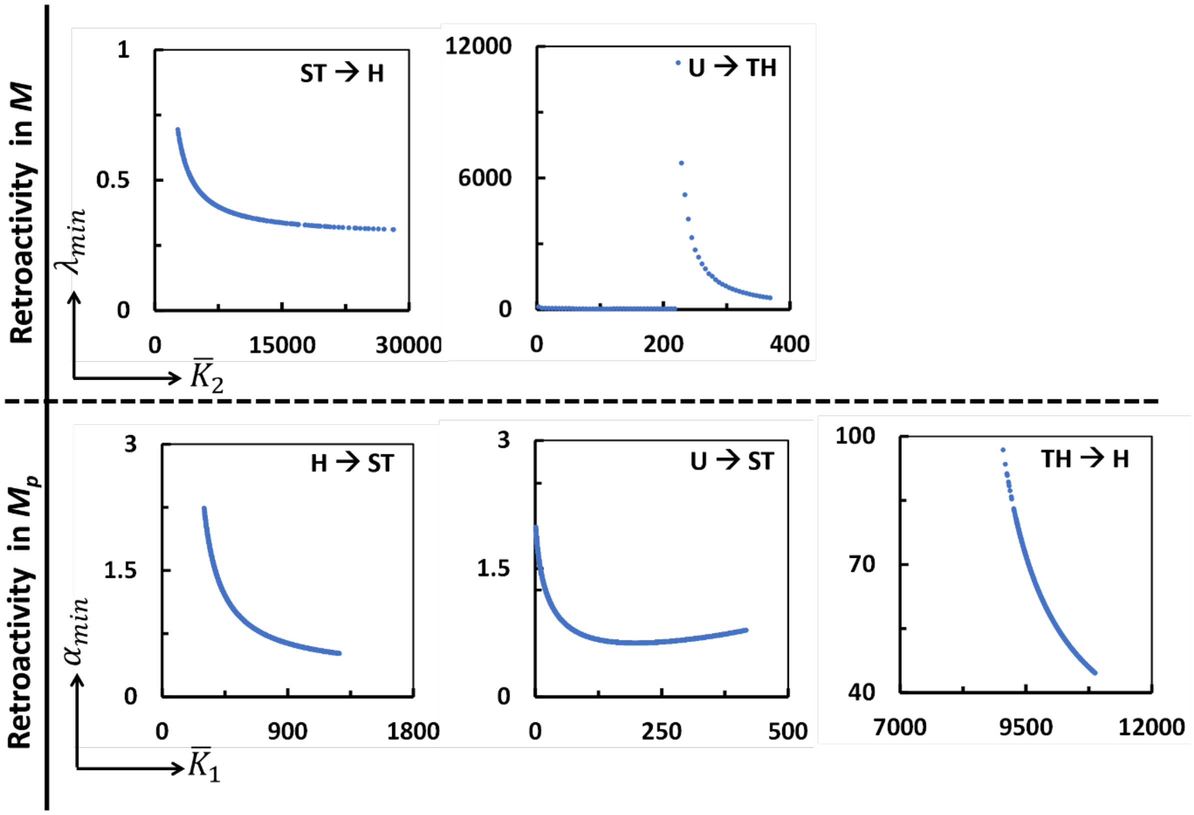


**Figure S2**: Effect of $\overline{K}_2(\alpha = 0)$ on $\lambda_{min}$ for the transitions (A) ST to H and (B) U to TH, and of $\overline{K}_1(\lambda = 0)$ on $\alpha_{min}$ for the transitions (C) H to ST, (D) U to ST, and (E) TH to H.

**S3. Sensitivity and rate balance analysis to investigate retroactivity induced regime transition.**

The retroactivity induced regime transition for ST to H (Fig S3), TH to H (FigS4) and H to ST (Fig S5) is investigated with the help of sensitivity of load ($\lambda$ $or$ $\alpha$) to enzyme concentration $e_t$ and rate balance analysis for different loads.

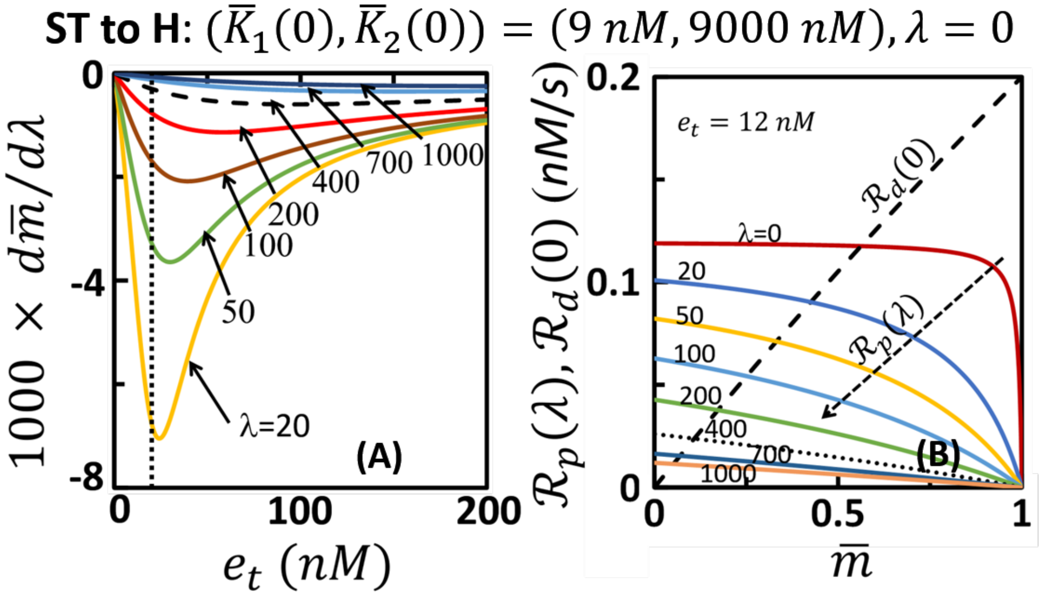


**Figure S3**: Sensitivity (A) and rate balance (B) plot consisting of the locus of $R_p(e_t = 12\ nM, \lambda, \bar{m})$ and $R_d(\alpha = 0\ , \bar{m})$ for ST to H regime transition.

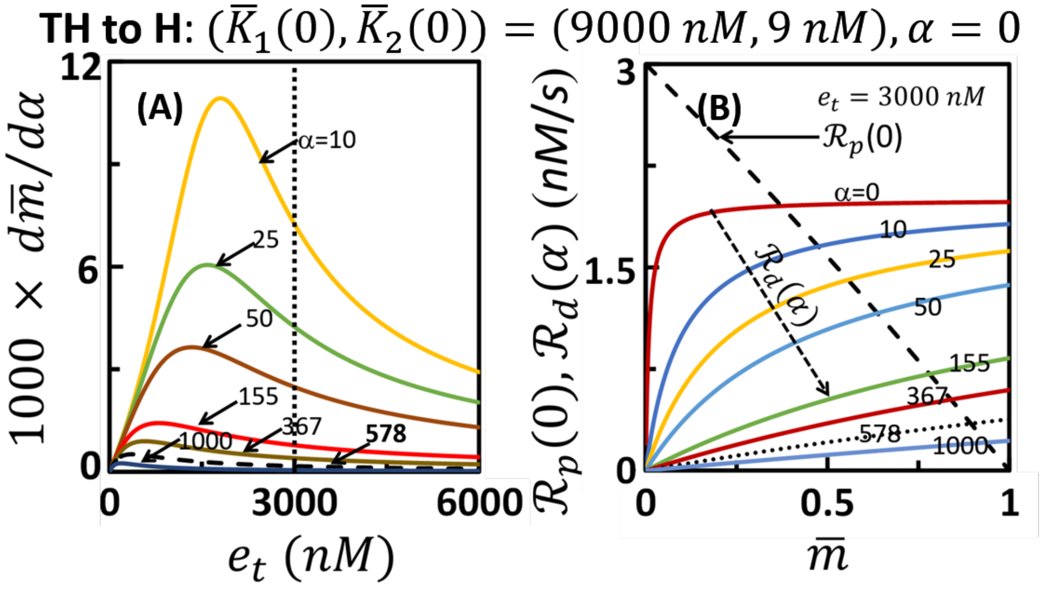


**Figure S4**: Sensitivity (A) and rate balance (B) plot consisting of the locus of $R_p(\lambda = 0, \bar{m})$ and $R_d(e_t = 3000\ nM, \alpha, \bar{m})$ for TH to H regime transition.

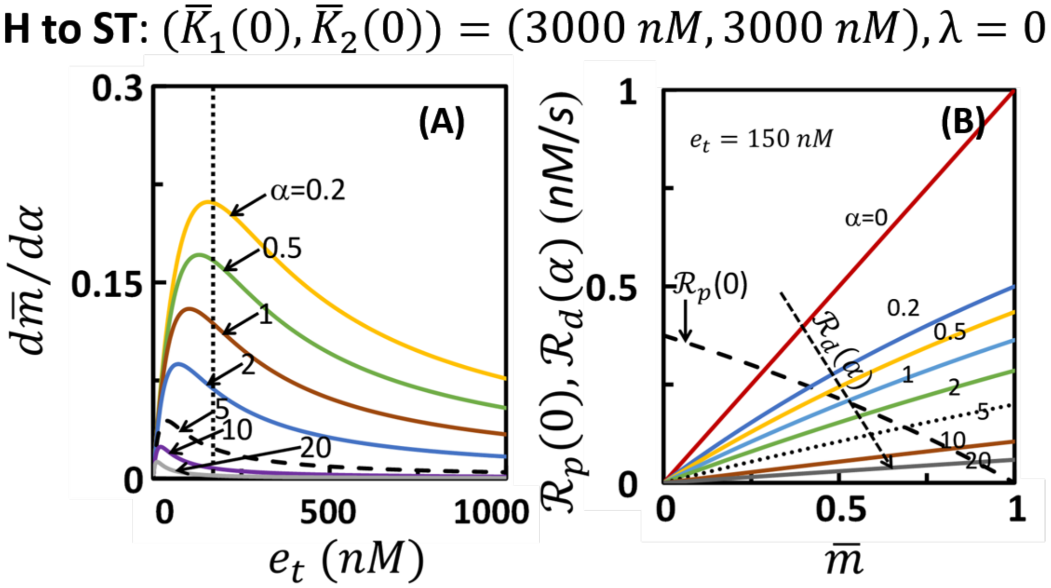


**Figure S5**: Sensitivity (A) and rate balance (B) plot consisting of the locus of $R_p(\lambda = 0, \bar{m})$ and $R_d(e_t = 150\ nM, \alpha, \bar{m})$ for H to ST regime transition.
